# *PURPL* identifies a poor outcome subgroup within Luminal A breast cancer

**DOI:** 10.1101/2025.11.11.687413

**Authors:** Salvador Junior Ochoa Zavalza, Maxime Pouokam, Masato Kruse, Georgina Gonzalez-Isunza, Radmila Sazdanovic, Javier Arsuaga

## Abstract

**Introduction:** Luminal A is the most common breast cancer molecular subtype and is generally associated with a favorable five-year prognosis. However, a substantial subset of patients experience relapse, suggesting that biologically distinct high-risk tumors exist within this subtype.

**Methods:** We applied Topological Analysis of Array Comparative Genomic Hybridization (TAaCGH) to identify recurrent copy number alterations associated with Luminal A breast cancer. We subsequently evaluated the long non-coding RNA *PURPL* using copy number, gene expression, and survival data in the independent TCGA, METABRIC, and SCAN-B cohorts. Associations with clinical outcome were examined using multivariable Cox regression and *TP53*-stratified analyses.

**Results:** *PURPL* copy number gain was associated with significantly worse survival in TCGA and METABRIC, while elevated *PURPL* expression was associated with significantly worse survival in TCGA and SCAN-B. Elevated *PURPL* expression remained associated with survival after adjustment for clinical covariates. In analyses stratified by *TP53* mutation status, high *PURPL* expression remained associated with worse survival in *TP53* wild-type tumors. Although *PURPL* has previously been implicated in cancer-related mechanisms, its copy number alteration, expression, and association with clinical outcome have not been systematically evaluated in Luminal A breast cancer.

**Conclusion:** These findings identify *PURPL* as a candidate lncRNA within a recurrent copy number alteration associated with adverse clinical outcome in Luminal A breast cancer and support further mechanistic investigation of its role in this subtype.

## Introduction

Breast cancer is the most commonly diagnosed cancer in women worldwide^1,2^. Luminal A, which accounts for more than 40% of all breast cancers^3^, is characterized by expression of the endocrine receptors estrogen and progesterone, lack of expression of the growth factor gene *ERBB2*, and low expression of the proliferation marker Ki67^4–6^. Luminal A tumors are commonly associated with good prognosis over five years, and in some cases, treatment does not include systemic chemotherapy^7^. However, despite this generally favorable prognosis, more than 25% of patients succumb to the disease later in life^8,9^. This suggests that Luminal A breast cancer is not a uniformly low-risk disease, but instead includes biologically distinct tumors with more aggressive clinical behavior^10,11^. Identifying molecular alterations associated with these high-risk tumors remains an important challenge.

Long non-coding RNAs (lncRNAs) are untranslated transcripts, typically several hundred nucleotides in length, that regulate diverse cellular processes through interactions with the genome, epigenome, other RNAs, and proteins^12–17^. Dysregulation of lncRNAs has been associated with multiple diseases^16,18–21^, including cancer^16,20,21^. In breast cancer, lncRNAs have been linked to aberrant cell cycle activity, metastasis progression, and chemotherapy resistance^22–29^. However, the role of many lncRNAs in defining clinically relevant subgroups within Luminal A tumors remains poorly understood^22,30^.

A frequent mechanism of gene expression dysregulation in cancer involves genome copy number changes^31–34^. Common copy number changes in Luminal A tumors include gain of chromosome arm 1p, loss of chromosome arm 16q, gains in chromosome arms 8q and 11q, and losses in chromosome arms 8p and 13q^3,35,36^. Although conventional statistical methods reliably identify the most recurrent copy number changes, alterations restricted to subsets of patients may be more difficult to detect consistently^36^. This is particularly relevant for Luminal A breast cancer, where clinically important high-risk tumors may represent only a subset of the subtype.

Topological data analysis provides a complementary framework for identifying heterogeneous genomic patterns by representing data as point clouds and using their topological structure to detect biologically meaningful differences among patient groups^37–42^. We previously introduced a topological approach for detecting copy number changes from array CGH data^43^. This method, called TAaCGH, performs statistical associations between phenotypes and topological summaries of patient copy number profiles. We later showed that descriptors such as *β*_0_ curves, lifespan curves^44^, and persistent landscapes^45^ identify copy number changes associated with cancer phenotypes, including a copy number alteration in chromosome region 5p14.3-p12 associated with Luminal A breast cancer^46^.

In the present study, we further investigated the chromosome region 5p14.3-p12 highlighted by TAaCGH and characterized the long non-coding RNA *PURPL*, also known as *LINC01021*, as a candidate transcript associated with poor outcome in Luminal A breast cancer. *PURPL* has been implicated in cancer-related processes across several tumor types, including colorectal cancer, lung cancer, melanoma, gastric cancer, thyroid cancer, and breast cancer^47–53^. In breast cancer, elevated *PURPL* expression has been linked to tumor-promoting immune effects and was reported to enhance macrophage recruitment and M2 polarization through *CCL2* signaling^54^. In addition, several studies have linked *PURPL* to p53-associated regulatory pathways. In colorectal cancer cells, *PURPL* was identified as a p53-induced lncRNA that suppresses basal p53 levels through interactions involving *MYBBP1A*^47^, while more recent work showed that loss of p53-induced *PURPL* expression enhances the induction of selected p53 target genes involved in apoptosis and stress responses^55^. These studies suggest that *PURPL* may influence both immune-related and p53-associated pathways. However, despite growing evidence for biological functions of *PURPL*, its clinical significance in Luminal A breast cancer remains poorly understood.

To evaluate its association with clinical outcome, we examined copy number, gene expression, and survival across multiple independent breast cancer cohorts. Across TCGA, METABRIC, and SCAN-B, *PURPL* copy number gain and elevated *PURPL* expression were consistently associated with poorer survival among patients with Luminal A breast cancer^33,56–58^. We further show that elevated *PURPL* expression remains associated with survival after adjustment for clinical covariates and that the survival difference associated with high *PURPL* expression is preserved in *TP53* wild-type tumors. Together, these findings identify *PURPL* as a candidate transcript associated with adverse clinical outcome in Luminal A breast cancer and support further investigation of its biological role, including its relationship with *TP53*-associated pathways in this subtype.

## Data and Methods

### Horlings Dataset

The Horlings dataset, initially presented in^31^, comprises 68 well curated breast cancer patient samples, including 21 classified as Luminal A. Bacterial artificial chromosome (BAC) microarray copy number profiles and molecular phenotype annotations were available for these samples. Because the copy number data are not segmented, this dataset is amenable to topological analysis of probe level copy number profiles.

### TCGA BRCA Dataset

We obtained data from the TCGA-BRCA project using the TCGABiolinks library (RRID:*SCR*_017683) in Bioconductor (RRID:*SCR*_006442)^59^. The dataset contains 540 Luminal A cases with available survival and genomic data, including copy number alterations, gene expression, and mutation status. Copy number alteration values were normalized and segmented to integer values in the range *{*1, 7*}*. The distribution of *PURPL* absolute copy number (ACN) values across patients is summarized in Supplementary Table S1. Because the sample sizes for ACN= 1 and ACN= 7 were small, subsequent analyses were restricted to samples with ACN values between 2 and 6. Gene expression data were normalized using the variance stabilizing transformation (VST) implemented in the DESeq2 package version 1.40.2 in R (RRID:*SCR*_015687)^60^. Mutation status was encoded as a binary variable. Survival data included each patient’s recorded survival time and event status, with follow-up times extending up to 8,000 days.

### METABRIC Dataset

The METABRIC cohort contains 700 Luminal A breast cancer patients^33,57^ and was accessed through cBioPortal for Cancer Genomics (RRID:*SCR*_014555)^61–63^. Of these patients, 449 had complete survival and copy number data for the genes or genomic regions analyzed in this study. Because the lncRNA *PURPL* was not directly represented in the METABRIC copy number annotation, copy number alterations were assessed at the *PURPL* genomic locus (chr5:27,217,714–27,497,871; hg38), using coordinates obtained from HGNC/Ensembl annotation. Copy number values were discretized into two categories: gain or amplification (GAIN), indicating a moderate to high increase in copy number, and neutral (NEUT), indicating no detectable change from baseline. Survival data included each patient’s recorded survival time and event status, with follow-up times extending up to 10,000 days.

### SCAN-B Dataset

The SCAN-B cohort, available through the Gene Expression Omnibus (GEO; RRID:*SCR*_005012) under accession GSE96058, contains RNA-seq data from 3,273 primary breast cancer patients^64,65^. For this study, we matched tumor samples with transcriptomic and mutation data, with mutation data obtained from the SCAN-B Mutation Explorer project (RRID:*SCR*_027895). The final analysis included 1, 609 Luminal A patients with both gene expression profiles and mutation status across key cancer associated genes^58^. Transcriptomic data were reported as Fragments Per Kilobase Million (FPKM) plus 0.1 and were log_2_ transformed. The values for *PURPL* ranged from −3.322 to 1.297. Survival data included each patient’s recorded survival time and event status, with follow-up times extending up to 2,500 days.

All patient data used in this study were anonymized prior to analysis. Investigators did not have access to personal identifiers.

### Topological Analysis of aCGH (TAaCGH)

TAaCGH uses a topological characterization of array CGH data to associate genomic regions with patient phenotypes^66^. An overview of the TAaCGH workflow is provided in the Results section. The methodological framework builds on earlier applications of computational homology to CGH profiles^43^, theoretical developments supporting the sliding-window topological representation of genomic data^67^, and the introduction of TAaCGH for identifying subtype-specific copy number aberrations^66^. Subsequent work extended the framework to predictive modeling and classification^36^, while more recent developments incorporated persistence curves and stability analyses within the TAaCGH suite^46^. Next, we describe the additional modules used to prioritize candidate genes from significant copy number regions.

Each significant region identified by TAaCGH contains approximately 20 probes, as illustrated in Figure 1c. For each probe within a significant region, we tested whether the average copy number value was significantly different from zero using a permutation test with 100,000 randomizations. We also recorded the sign of the mean copy number value to determine whether the region showed a gain or a loss (Figure 1c). To determine whether the selected probes shows a significant copy number difference in the test set relative to the control set, we compared average copy number values between groups. All p-values were adjusted for the number of probes in each region using the Benjamini-Hochberg (BH) false discovery rate correction^68^.

**Figure 1.**
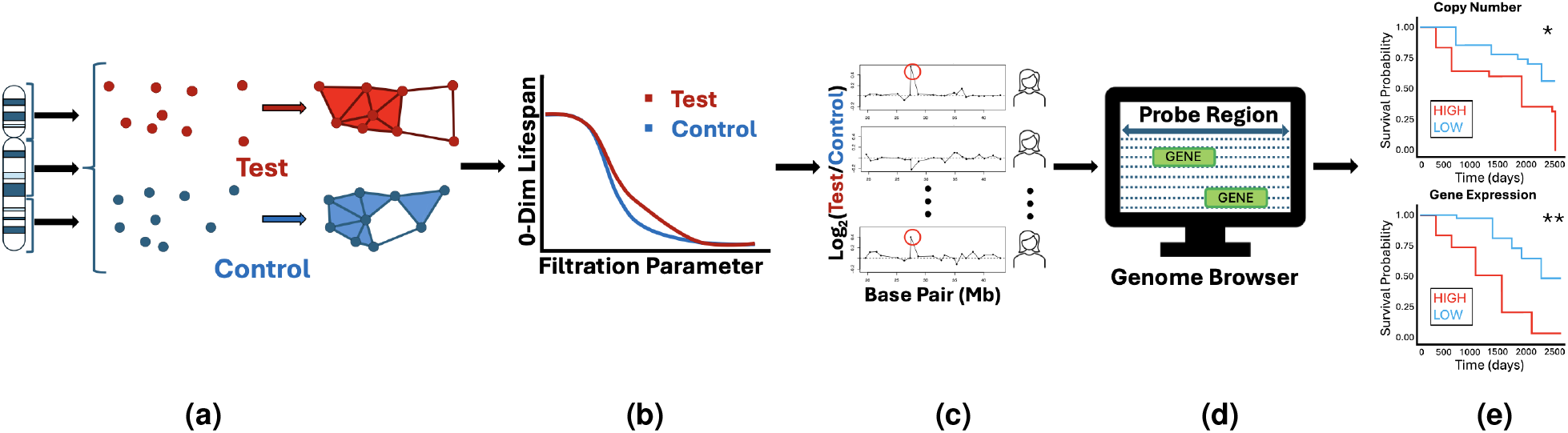
TAaCGH workflow. The figure summarizes the steps used to identify genes associated with a given phenotype and their clinical implications: (a) Copy number profiles are subdivided into fragments and patient samples are divided into test and control groups. A point cloud, and its corresponding filtrations, are generated for each fragment and patient group. (b) Persistent homology computation of topological descriptors determine copy number changes associated to the test set. (c) Probe level analysis of significant regions using BH-corrected statistical testing; (d) Retrieval of nearby genes from a Genome Browser. (e) Survival analysis and validation using TCGA/METABRIC/SCAN-B datasets.

For each significant probe, the associated genomic interval was defined as the region bounded by the two immediately adjacent probes. This interval was queried in the UCSC Genome Browser (RRID:*SCR*_005780) to identify genes located in the associated genomic interval. Candidate genes identified in this step were then evaluated using copy number, gene expression, survival, and mutation data from independent cohorts.

### Survival Analysis

To ensure comparable follow-up across cohorts, survival times were administratively censored before analysis. For copy number analyses, survival times in TCGA and METABRIC were censored at 8,000 days, corresponding to the maximum follow-up time available in TCGA. For gene expression analyses, survival times in TCGA and SCAN-B were censored at 2,500 days, corresponding to the maximum follow-up time available in SCAN-B.

Survival differences were evaluated using a two step analytical process. First, patients in each cohort were divided into high and low groups according to the Maximally Selected Rank Statistic (MaxStat) test^69,70^. MaxStat identifies the cutoff point *µ* that best dichotomizes the patient population by copy number or gene expression values, defined as the value that maximizes the absolute log-rank statistic: *M* = max_*µ*_ |*S*_*µ*_| Additionally, MaxStat provides a p-value associated with the maximally selected log-rank statistic. Throughout the text, we report the values of *µ* and *M*, while the corresponding MaxStat p-values are provided in Supplementary Tables S3 and S4. Kaplan-Meier curves were then used to visualize survival differences between the high and low groups defined using *µ*. The p-values reported in the main text correspond to log-rank tests comparing survival between these groups. When multiple related survival tests were performed, p-values were adjusted using the BH method.

### Cox Proportional Hazards Models

Cox proportional hazards models were used to test whether *PURPL* copy number or expression remained associated with overall survival after adjustment for relevant clinical covariates. The Cox model is semiparametric and estimates covariate effects without assuming a specific distribution for the baseline hazard over time.

Clinical factors associated with breast cancer outcomes, including age, tumor size, tumor grade, endocrine receptor status, and lymph node status, are commonly used in Cox models^71,72^. To reduce the risk of overfitting and keep the models interpretable, we limited model complexity in the TCGA and SCAN-B analyses to *PURPL*, age, tumor size, and lymph node status. The proportional hazards assumption was evaluated using Schoenfeld residuals as implemented in the cox.zph function of the survival package version 3.5-5 in R (RRID:*SCR*_021137)^73,74^. Variables that violated this assumption were included as stratification factors when appropriate.

To assess whether the association between *PURPL* expression and survival varied by *TP53* mutation status, Cox proportional hazards models were fitted separately in *TP53* wild-type and *TP53* mutant tumors when sample size permitted. For subgroups with limited sample sizes or sparse events, Firth-penalized and Bayesian Cox proportional hazards models were used as sensitivity analyses. Firth penalization was applied to address separation and unstable estimates, while the Bayesian Cox model used a weakly informative prior on the log hazard ratio to stabilize estimation.

## Results

### TAaCGH detects a chromosome gain of lncRNA *PURPL* in Luminal A patients

Copy number alterations associated with Luminal A breast cancer were identified using TAaCGH, a statistical genetics method that associates patient phenotypes with topological representations of copy number profiles. Using a sliding window algorithm, each segment of the genome is assigned a two dimensional point cloud and the corresponding simplicial complex is constructed (Figure 1a). Functional topological descriptors, including lifespan and landscape curves, are then used to identify genomic regions that warrant further analysis^66,67^ (Figure 1b).

In this work, we extend TAaCGH by testing the significance of individual probes within previously detected regions (Figure 1c), and by identifying cancer related genes located near significant probes using the UCSC Genome Browser, (Figure 1d). Candidate genes were then evaluated using copy number, gene expression, and survival data. Patients were divided into high and low groups according to copy number or gene expression values using the Maximally Selected Rank Statistic (MaxStat) test, and survival differences were assessed using Kaplan-Meier and Cox Proportional Hazards survival analysis. Reproducibility was evaluated across TCGA, METABRIC, and SCAN-B (Figure 1e).

The chromosomal region 5p14.3–p12 was previously classified as significant by 0-dimensional lifespan Betti curves and third persistent landscape curves^46,75^ (Figure 2). Figure 2a shows the lifespan Betti curve for Luminal A patients in red and control patients in blue. The slower decrease in the average 0-th Betti numbers in Luminal A patients indicates a difference in the topological structure of copy number profiles between the two groups. Differences between the groups remained significant after multiple testing correction, with a BH adjusted p-value of *p* = 1.0 *×* 10^−5^ for lifespan curves and *p* = 0 for the second and third topological landscapes (Figure 2b); see^46^ for a detailed analysis.

**Figure 2.**
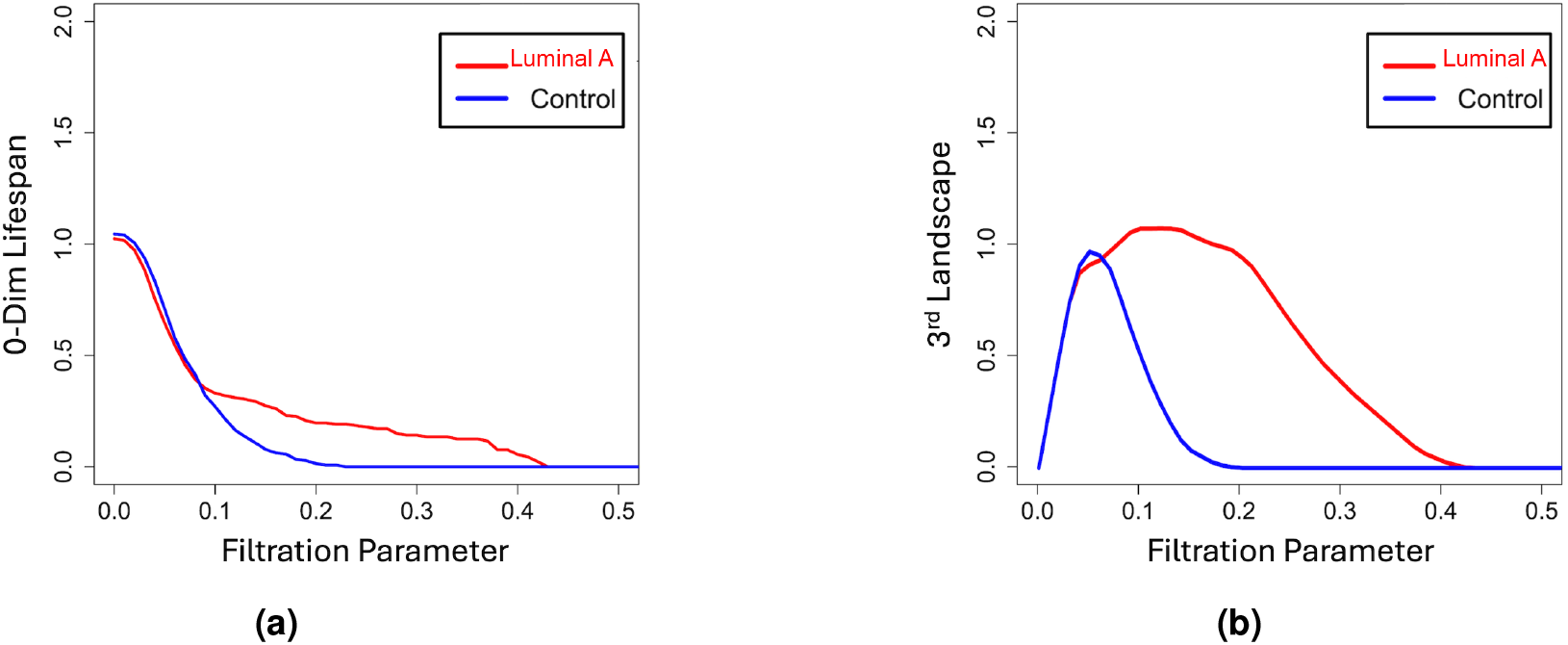
Topological descriptors of copy number for chromosome region 5p14.3-p12 for Luminal A and control patients: average (a) lifespan curves, and (b) 3rd landscape curves.

This region contains 21 probes and extends from basepair position 19, 608, 109 to 42, 861, 888. Probe level analysis showed that the average copy number values for probes CTD-2291F22 and RP11-46C20 differed significantly between Luminal A and control patients. However, only RP11-46C20 was driven by amplification in Luminal A patients, with a BH adjusted value of *p* = 0.016. Figure 3 shows two Luminal A patient profiles with a visually detectable copy number change at probe RP11-46C20, located at basepair position 27,452,428. Additional patient profiles are shown in Supplementary Figure S1.

**Figure 3.**
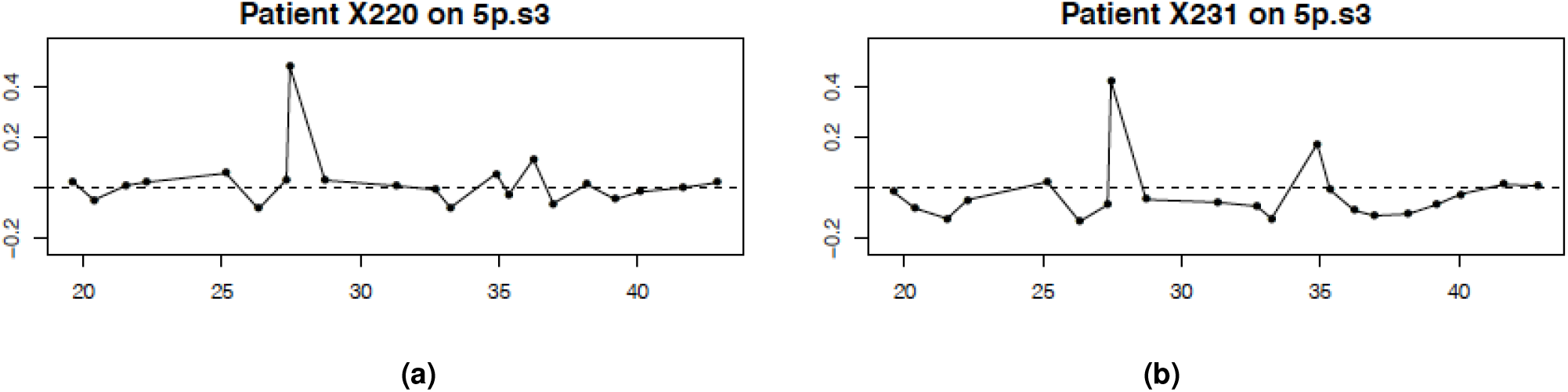
Luminal A patient profiles from the Horlings dataset on cytobands 5p14.3-p12. The x-axis shows the genomic position (Mb) and the y-axis *log*_2_ fold-change in mean copy number values comparing Luminal A samples to the control group. The profiles show an amplification of the *RP*11-46*C*20 at position 27,452,428.

The UCSC Genome Browser^76,77^ identified a single gene located at probe RP11-46C20: *PURPL*. We also evaluated the associated genomic region comprised of CTD-2219P12 and RP11-37M16, located at basepair positions 27,325,194 and 28,716,023, respectively. In addition to *PURPL*, this search identified *LINC02103* and the pseudogene *PNU6-909P*. Since *PURPL* has been associated with tumor processes in other cancers^47–53^, we hypothesized that a subset of Luminal A tumors is characterized by amplification of *PURPL*.

### *PURPL* copy number changes are associated with patient survival in Luminal A patients

To test whether *PURPL* amplification was associated with survival in Luminal A breast cancer, we first analyzed the TCGA dataset. The MaxStat test identified an ACN cutoff of *µ* = 3 (*M* = 2.15), dividing patients into high ACN (> 3) and low ACN (*≤* 3) groups. Kaplan-Meier survival analysis using this cutoff showed a significant survival difference, with *p* = 0.016 (Figure 4a). Luminal A patients with high *PURPL* ACN had significantly worse survival than patients with low *PURPL* ACN.

**Figure 4.**
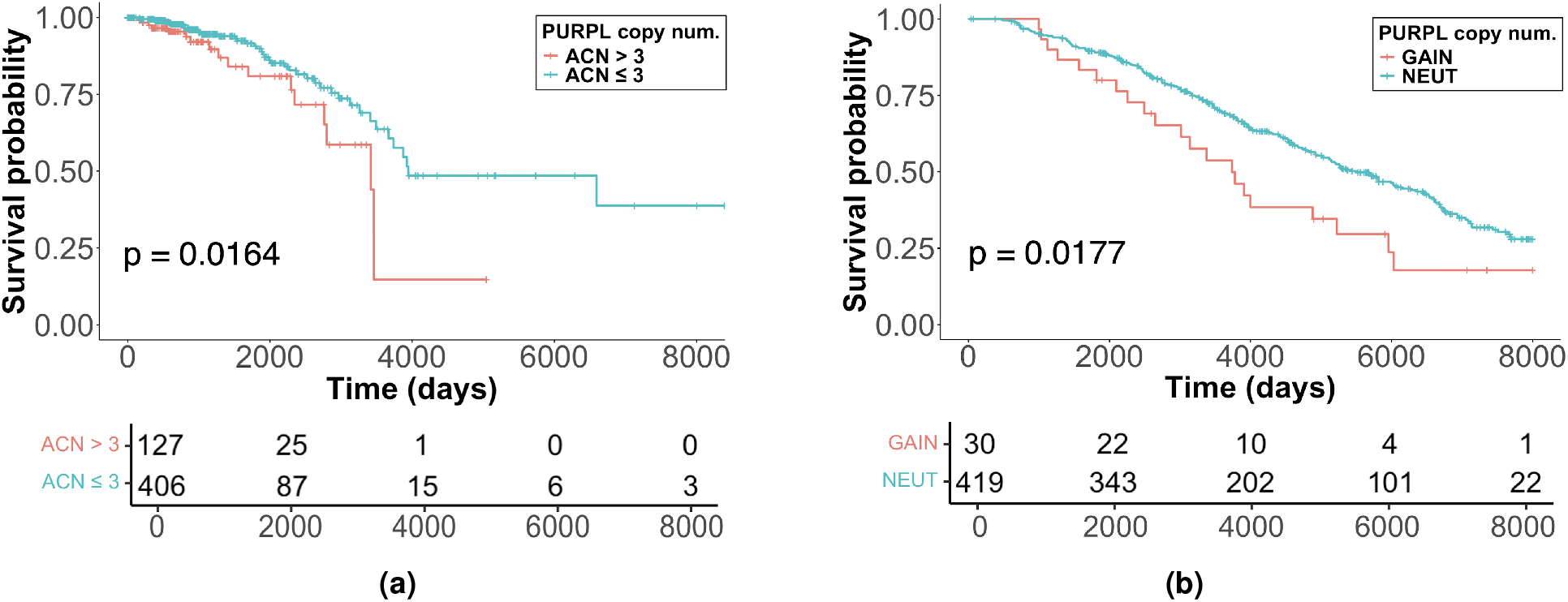
*PURPL* copy number survival analysis: Kaplan-Meier curves stratified by (a) high vs. low *PURPL* ACN in TCGA and (b) *PURPL* GAIN vs. NEUT in METABRIC.

Next we sought to determine whether this association could be replicated in the METABRIC dataset. Among 449 Luminal A patients, 30 had *PURPL*-GAIN, while the remaining patients were classified as *PURPL*-NEUT. Survival analysis using these preset copy number categories showed a significant survival difference (*p* = 0.018; Figure 4b), with *PURPL*-GAIN patients showing poorer survival than patients in the NEUT category. This pattern was consistent with the TCGA copy number analysis, in which increased *PURPL* copy number was also associated with reduced survival. The METABRIC findings support an association between elevated *PURPL* copy number and adverse clinical outcome in Luminal A breast cancer.

### Overexpression of *PURPL* is associated with patient survival in Luminal A patients

Because copy number gain can alter gene expression, we then examined whether elevated *PURPL* expression was also associated with survival in Luminal A patients. In TCGA, the MaxStat procedure identified an optimal gene expression cutoff of *µ* = 4.90 (*M* = 2.76). Kaplan-Meier survival analysis using this cutoff showed a significant survival difference (*p* = 7.35 *×* 10^−4^), with high *PURPL* expression associated with worse prognosis (Figure 5a).

**Figure 5.**
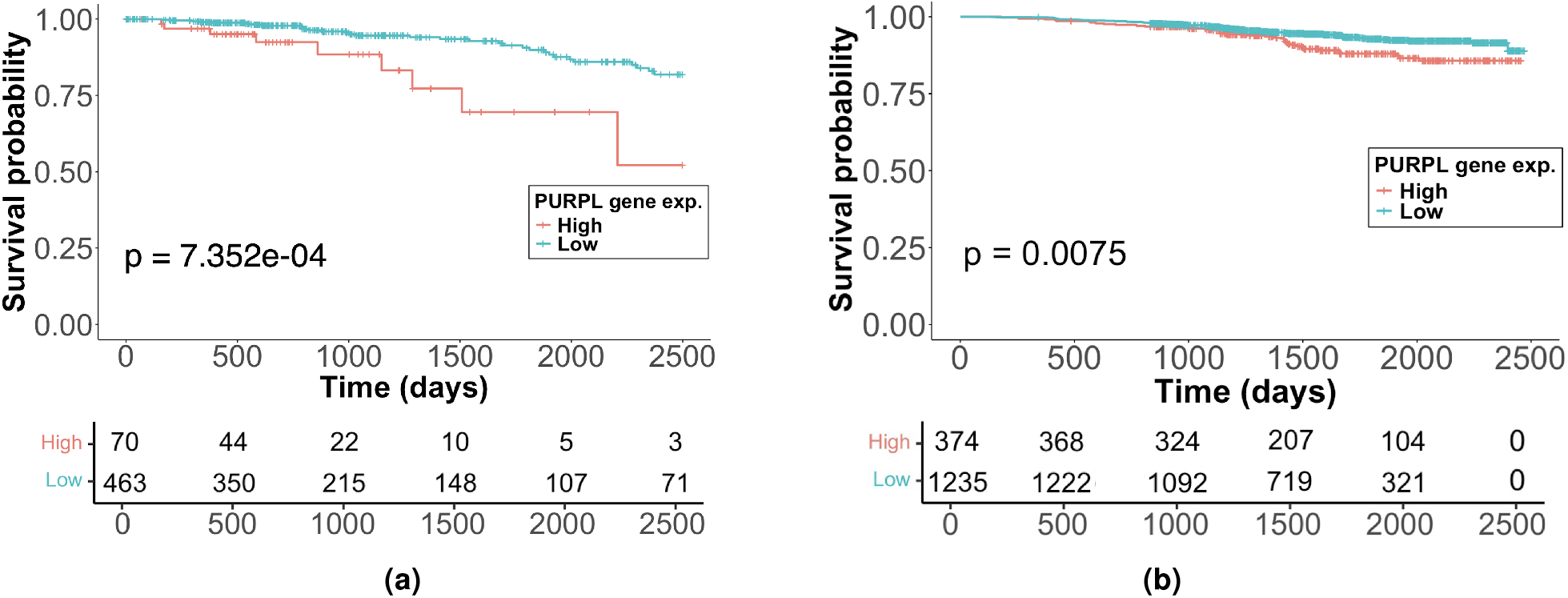
*PURPL* gene expression survival analysis: Kaplan-Meier curves stratified by high vs. low *PURPL* expression for (a) TCGA and (b) SCAN-B data.

Since METABRIC does not contain expression data for *PURPL*, we evaluated *PURPL* expression in SCAN-B. The MaxStat cutoff was *µ* = −2.80 (*M* = 2.66), assigning 374 patients to the high-expression category. High *PURPL* expression was associated with worse survival (*p* = 0.0075) (Figure 5b).

Additional analyses were performed to determine whether the association between *PURPL* expression and survival extended to other breast cancer subtypes (Table 1). Although high *PURPL* expression was significantly associated with survival in the Luminal B and HER2 enriched subtype in TCGA (*p*_*LumB*_ = 0.0093, *n*_*LumB*_ = 195; *p*_*HER*2_ = 0.025, *n*_*HER*2_ = 78), this result was not validated in SCAN-B (*p*_*LumB*_ = 0.21;*p*_*HER*2_ = 0.21). No other subtype was significant. These results indicate that the association between *PURPL* overexpression and worse survival is most consistently observed in Luminal A tumors.

**Table 1.**
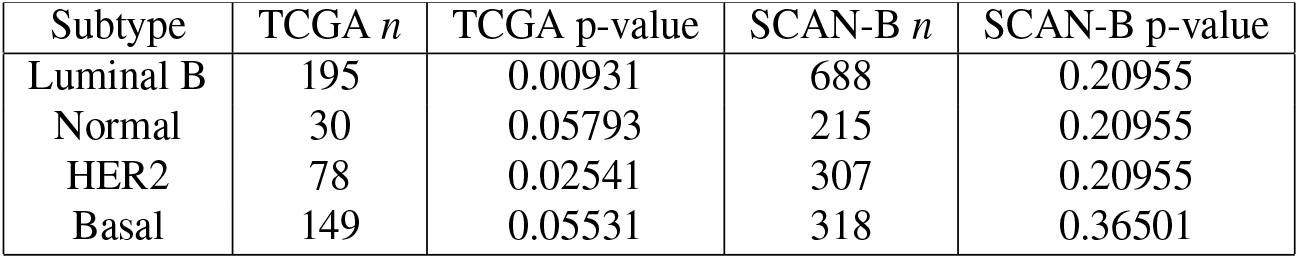
Association between *PURPL* expression and overall survival across additional breast cancer subtypes in the TCGA and SCAN-B datasets.

| Subtype | TCGA $n$ | TCGA p-value | SCAN-B $n$ | SCAN-B p-value |
| --- | --- | --- | --- | --- |
| Luminal B | 195 | 0.00931 | 688 | 0.20955 |
| Normal | 30 | 0.05793 | 215 | 0.20955 |
| HER2 | 78 | 0.02541 | 307 | 0.20955 |
| Basal | 149 | 0.05531 | 318 | 0.36501 |

### *PURPL* expression and *TP53* mutation status

Previous studies have linked *PURPL* to p53-associated regulatory pathways. In colorectal cancer cells, *PURPL* was identified as a p53-induced lncRNA that suppresses basal p53 levels through interactions involving *MYBBP1A*, thereby promoting tumor cell growth^47^. In liver cancer, *PURPL* is overexpressed in tumor tissues and cell lines, and experimental knockdown of *PURPL* increases p53 expression while reducing cell proliferation^48^. More recently, Kaller and colleagues showed that loss of p53-induced *PURPL* expression enhances the induction of selected p53 target genes involved in apoptosis and stress responses, including *NOXA* and *FAS*, and alters transcript splicing patterns, suggesting that *PURPL* normally attenuates specific components of the p53 regulatory network^55^. These observations suggest that *PURPL* participates in p53-associated regulatory pathways. Because several reported functions of *PURPL* depend on intact p53 signaling, we examined whether the association between *PURPL* expression and survival differed between *TP53* wild-type and *TP53* mutant tumors.

TCGA Luminal A patients were divided into tumors with no detected *TP53* mutation and tumors with a detected *TP53* mutation, denoted as *TP*53_*wt*_ and *TP*53_*mut*_, respectively. In the *TP*53_*wt*_ group 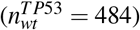, the MaxStat gene expression cutoff was *µ* = 4.90 (*M* = 3.03), and high *PURPL* expression was associated with worse survival, with a BH adjusted p-value of 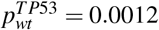. We validated this finding in SCAN-B. In the SCAN-B *TP*53_*wt*_ group 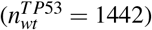, the MaxStat cutoff was *µ* = −2.80 (*M* = 2.66), and high *PURPL* expression was again associated with worse survival, with a BH adjusted p-value of 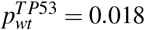. Kaplan-Meier curves are shown in Figure 6. Notably, the association between high *PURPL* expression and adverse survival outcomes persisted in Luminal A tumors without detectable *TP53* mutations.

**Figure 6.**
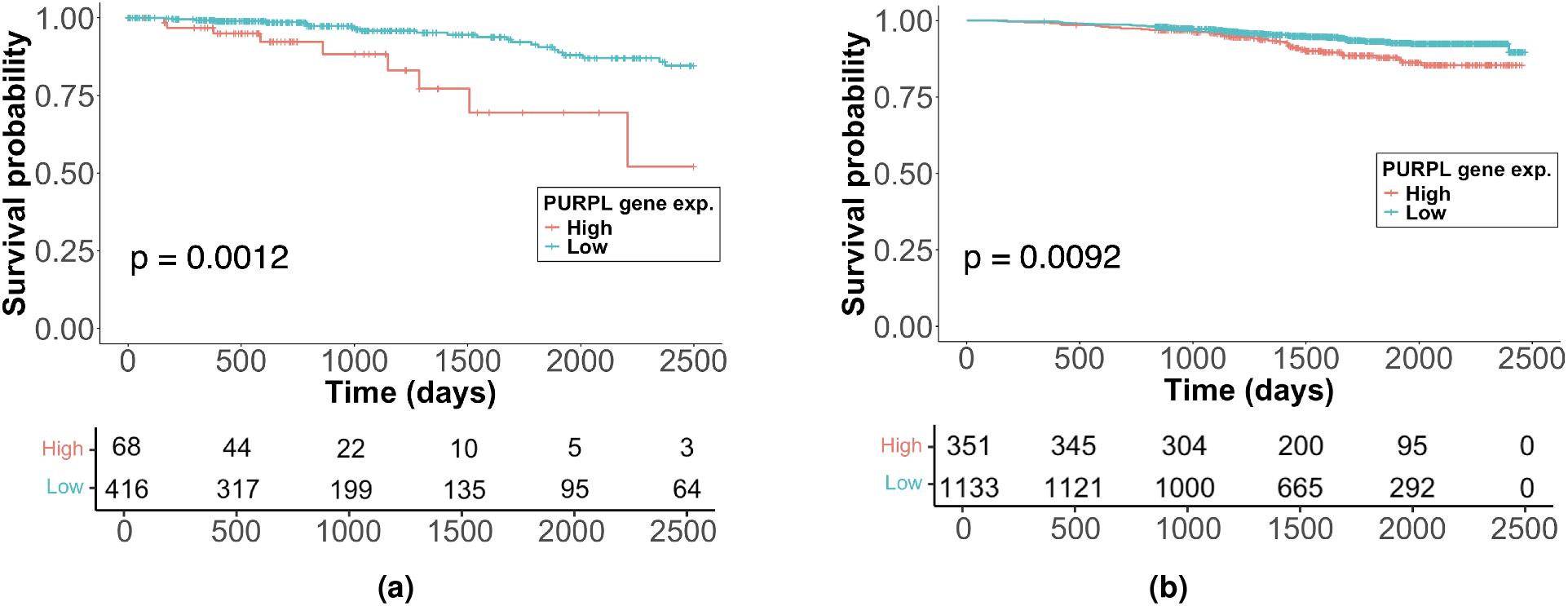
*TP*53_*wt*_ survival analysis: Kaplan-Meier curves stratified by high vs. low *PURPL* expression among patients with *TP*53_*wt*_ for (a) TCGA and (b) SCAN-B.

In TCGA, the *TP*53_*mut*_ subgroup included 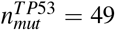 patients. The MaxStat gene expression cutoff was *µ* = 4.10 (*M* = 1.94), with a BH adjusted p-value of 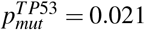. In SCAN-B, the *TP*53_*mut*_ subgroup included 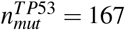 patients. The MaxStat cutoff was *µ* = −2.48 (*M* = 1.31), with a BH adjusted p-value of 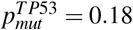, see Supplementary Tables S3-S7 for detailed results. Thus, the survival association of *PURPL* expression was most consistently observed in *TP*53_*wt*_ Luminal A tumors and was not consistently validated in *TP*53_*mut*_ tumors.

### *PURPL* remains associated with survival after adjustment for clinical covariates

In the Kaplan-Meier analyses, *PURPL* copy number and gene expression were associated with overall survival in Luminal A breast cancer. We therefore used Cox proportional hazards models to test whether these associations remained after adjustment for clinically relevant covariates.

Age, lymph node status, and tumor size were selected as clinically relevant covariates for breast cancer survival analysis. Age at diagnosis has been shown to influence breast cancer prognosis and survival outcomes across molecular subtypes^78^. Lymph node status has been associated with clinical outcomes in breast cancer and across molecular subtypes^71,72^. Tumor size is an established prognostic factor and has been associated with breast cancer survival independent of nodal involvement^79^. These covariates were therefore included in the multivariable Cox proportional hazards models.

Similar to the Kaplan-Meier analysis, the Cox models were evaluated in two datasets: TCGA and SCAN-B. In TCGA, after removing entries with missing or non-assessable values, the final cohort included *n* = 461 patients. Age was split using the TCGA median age of 61.1 years. Lymph node status was classified as node-negative (*N*^−^; N0) or node-positive (*N*^+^; N1-N3). Tumor size was categorized as T1 (*≤*20 mm) versus T2-T4 (> 20 mm) and was used as a stratification factor when the proportional hazards assumption was violated, allowing separate baseline hazard functions for each tumor size group while preserving estimation of the remaining covariate effects.

In SCAN-B, after removing entries with missing or non-assessable values, the final Luminal A cohort included *n* = 1565 patients. Age cutoff was also the median in this case (65 years). Lymph node status was classified as node-negative (*N*^−^) or node-positive (*N*^+^), and tumor size was categorized as *≤* 20 mm versus > 20 mm. Tumor size was included as a stratification factor when the proportional hazards assumption was violated.

In TCGA, the Cox model using *PURPL* copy number showed that high *PURPL* copy number was significantly associated with worse overall survival. Patients with *PURPL* copy number > 3 had a higher hazard of death compared with patients with copy number *≤* 3 (HR = 2.34, 95% CI: 1.18–4.62, *p* = 0.0146). Age was also significantly associated with worse survival (HR = 2.70, 95% CI: 1.33–5.49, *p* = 0.0060), whereas lymph node status did not reach statistical significance (Table 2).

**Table 2.** Cox model using *PURPL* copy number (> 3 vs *≤* 3), age with median split, and node positive vs node negative covariates in TCGA.

| Variable | Hazard Ratio (HR) | 95% CI | p-value |
| --- | --- | --- | --- |
| *PURPL* copy number $> 3$ vs $\leq 3$ | 2.34 | 1.18–4.62 | 0.0146 |
| Age $> 61.1$ vs $\leq 61.1$ years | 2.70 | 1.33–5.49 | 0.0060 |
| $N^+$ vs $N^-$ | 1.90 | 0.95–3.81 | 0.0713 |

The TCGA Cox model was also fitted using *PURPL* gene expression (Table 3). High and low expression groups were defined using a MaxStat-derived split point of 4.90. The same covariates were used as in the copy-number model. High *PURPL* expression was significantly associated with worse overall survival (HR = 4.20, 95% CI: 1.74–10.13, *p* = 0.0014). Age also remained significantly associated with worse survival (HR = 3.21, 95% CI: 1.59–6.47, *p* = 0.0011), while lymph node status was not statistically significant.

**Table 3.** Cox model using *PURPL* high vs low expression (*µ* = 4.90), age with median split, and node positive vs node negative covariates in TCGA.

| Variable | Hazard Ratio (HR) | 95% CI | p-value |
| --- | --- | --- | --- |
| *PURPL* expression high vs low | 4.20 | 1.74–10.13 | 0.0014 |
| Age $> 61.1$ vs $\leq 61.1$ years | 3.21 | 1.59–6.47 | 0.0011 |
| $N^+$ vs $N^-$ | 1.86 | 0.91–3.80 | 0.0870 |

The Cox model was then fitted separately in patients with *TP*53_*wt*_ and *TP*53_*mut*_ tumors. High *PURPL* expression remained significantly associated with worse overall survival in the full Luminal A cohort (*p* = 0.0014) and in the *TP*53_*wt*_ subgroup (Table 4). In the *TP*53_*mut*_ subgroup, the standard Cox model was not estimable because no events occurred among patients with high *PURPL* expression. Sensitivity analyses using Firth penalized and Bayesian Cox models produced highly uncertain estimates and did not provide evidence for an association between *PURPL* expression and survival in the *TP*53_*mut*_ subgroup (Supplementary Tables S8 and S9).

**Table 4.** Cox model fitted for *TP*53_*wt*_ and *TP*53_*mut*_ patient subgroups using *PURPL* high vs low expression (*µ* = 4.90), age with median split, and node positive vs node negative covariates in TCGA.

| Variable | n | Hazard Ratio (HR) | 95% CI | p-value |
| --- | --- | --- | --- | --- |
| All patients: *PURPL* expression high vs low | 461 | 4.20 | 1.74–10.13 | 0.0014 |
| *TP53*<sub>wt</sub>: *PURPL* expression high vs low | 416 | 4.53 | 1.84–11.15 | 0.0010 |
| *TP53*<sub>mut</sub>: *PURPL* expression high vs low | 45 | Not estimable | - | - |

The Cox model was also fitted for the SCAN-B dataset using *PURPL* gene expression (Table 5). High and low expression groups were defined using a MaxStat split point of −2.80. The model used the same covariates as in the TCGA analysis, with tumor size included as a stratification variable because it did not satisfy the proportional hazards assumption. High *PURPL* expression was significantly associated with worse overall survival (HR = 1.51, 95% CI: 1.01–2.27, *p* = 0.0444). Age also remained significantly associated with worse survival (HR = 4.18, 95% CI: 2.63–6.64, *p* = 1.57 *×* 10^−9^), while lymph node status was not statistically significant.

**Table 5.** Cox model using *PURPL* high vs low expression (*µ* = −2.80), age with median split, and node positive vs node negative covariates in SCAN-B.

| Variable | Hazard Ratio (HR) | 95% CI | p-value |
| --- | --- | --- | --- |
| *PURPL* expression high vs low | 1.51 | 1.01–2.27 | 0.0444 |
| Age > 65 vs $\leq 65$ years | 4.18 | 2.63–6.64 | $1.569 \times 10^{-9}$ |
| $N^+$ vs $N^-$ | 0.97 | 0.64–1.47 | 0.8942 |

Consistent with the TCGA findings, high *PURPL* expression was associated with worse overall survival in the full SCAN-B Luminal A cohort (*p* = 0.0444) and in the *TP*53_*wt*_ subgroup (*p* = 0.0277), but not in *TP*53_*mut*_ tumors (Table 6). These results indicate that the association between elevated *PURPL* expression and poorer survival is primarily observed in *TP*53_*wt*_ tumors and is consistent across independent Luminal A breast cancer cohorts.

**Table 6.** Cox model fitted for *TP*53_*wt*_ and *TP*53_*mut*_ patient subgroups using *PURPL* high vs low expression (*µ* = −2.80), age with median split, and node positive vs node negative covariates in SCAN-B.

| Variable | n | Hazard Ratio (HR) | 95% CI | p-value |
| --- | --- | --- | --- | --- |
| All patients: *PURPL* expression high vs low | 1565 | 1.51 | 1.01–2.27 | 0.0444 |
| *TP53*<sub>wt</sub>: *PURPL* expression high vs low | 1404 | 1.62 | 1.05–2.47 | 0.0277 |
| *TP53*<sub>mut</sub>: *PURPL* expression high vs low | 161 | 0.93 | 0.25–3.49 | 0.9201 |

## Discussion

This study identifies the lncRNA *PURPL* as a transcript associated with poor outcome in Luminal A breast cancer. Although Luminal A tumors are generally associated with favorable prognosis, a substantial subset of patients experience relapse and die from disease. Starting from a recurrent copy number alteration in chromosome region 5p14.3–p12 identified by TAaCGH, we localized *PURPL* as a candidate transcript within this region and found that both *PURPL* copy number gain and increased *PURPL* expression were associated with worse survival. These findings identify *PURPL* as a candidate gene within a recurrent copy number alteration associated with poorer clinical outcome in Luminal A breast cancer and support further investigation of its biological role in this subtype.

The association between *PURPL* and clinical outcome was supported across independent datasets. *PURPL* copy number gain was associated with worse survival in both TCGA and METABRIC, while elevated *PURPL* expression was associated with worse survival in TCGA and SCAN-B. Importantly, multivariable Cox proportional hazards models showed that this association remained significant after adjustment for age, lymph node status, and tumor size. The concordance between copy number and expression analyses strengthens the evidence linking *PURPL* to poor clinical outcome.

An important aspect of this study is the strategy used to identify *PURPL*. Rather than focusing exclusively on the most recurrent genomic alterations across all Luminal A tumors, topology-guided analysis highlighted a recurrent copy number alteration present in a subset of Luminal A tumors. Subsequent analyses connected this topological signal to *PURPL* and linked its dysregulation to poor prognosis. These results illustrate how topological approaches can complement conventional genomic analyses by identifying alterations that may be clinically relevant despite occurring in only a fraction of tumors. Rather than replacing established genomic approaches, such methods may help prioritize candidate genomic regions and transcripts for further investigation.

Long non-coding RNAs are increasingly recognized as important contributors to breast cancer biology and clinical outcome. Prior studies have linked lncRNAs to tumor progression, therapy response, and prognosis, with some also implicating lncRNAs in p53-associated regulatory programs in breast cancer^22–29,80,81^. *PURPL* adds to the growing evidence that lncRNAs can provide prognostic information within breast cancer subtypes. In Luminal A disease, altered *PURPL* levels are associated with less favorable outcomes, supporting further evaluation of lncRNA-based markers in this subtype.

Several reported functions of *PURPL* provide biological plausibility for its association with poor prognosis. In breast cancer, *PURPL* has been linked to macrophage recruitment and M2 polarization through *CCL2* signaling, suggesting that it may contribute to a tumor-promoting immune microenvironment^54^. Similar immune-related functions have also been reported in lung cancer, where *PURPL* promotes M2 macrophage polarization through the RBM4/xCT signaling pathway^51^, indicating that immune modulation may represent a broader feature of *PURPL* biology. Beyond immune regulation, recent work has implicated *PURPL* in p53-dependent regulation of gene expression and alternative splicing. In colorectal cancer models, loss of p53-induced *PURPL* (*LINC01021*) altered the expression and splicing of several p53-responsive genes, including *LYN*^55^. Although these findings have not been examined in breast cancer, they are noteworthy because the full-length LYNA isoform promotes migration and invasion, and a high LYNA::LYNB ratio has been associated with shorter breast cancer-specific survival^82^. Collectively, these observations suggest that *PURPL* may influence tumor progression through multiple biological processes, including immune remodeling and regulation of gene expression.

The relationship between *PURPL* and p53 signaling is of particular interest. Previous studies have shown that *PURPL* participates in p53-associated regulatory networks, including suppression of basal p53 levels through interactions involving *MYBBP1A* and modulation of downstream p53-responsive genes^47,55^. In the present study, elevated *PURPL* expression remained associated with poorer survival in *TP53* wild-type tumors in both TCGA and SCAN-B, whereas evidence for a similar association in *TP53* mutant tumors was limited and inconsistent across cohorts. Although these findings do not establish a causal mechanism, they raise the possibility that the prognostic significance of *PURPL* is most evident in *TP53* wild-type tumors. In light of previous reports linking *PURPL* to p53 signaling and alternative splicing, it will be important to determine whether the clinical associations observed in Luminal A breast cancer reflect effects on p53-regulated transcription, RNA processing, immune remodeling, or a combination of these mechanisms.

Several limitations should be considered when interpreting these results. The analyses were based on retrospective cohorts, so prospective validation will be needed before *PURPL* can be evaluated for clinical risk stratification. In addition, the available datasets do not allow us to determine whether *PURPL* directly affects p53 activity or other tumor-related pathways. Functional data for genes previously linked to *PURPL*, including *MYBBP1A*, were also limited or unavailable. Finally, differences among cohorts in copy number annotation, expression measurement, clinical annotation, and follow-up time limit direct comparisons of effect sizes.

These findings support further investigation of *PURPL* in Luminal A breast cancer. Because Luminal A is the most common breast cancer molecular subtype, improved understanding of the molecular alterations underlying clinical heterogeneity within this group could have important biological and clinical implications. More broadly, this study illustrates how TDA can prioritize previously underexplored genomic regions for downstream investigation. By identifying alterations present within subsets of tumors rather than only the most recurrent genomic events, this strategy may help uncover candidate genes associated with the molecular heterogeneity of breast cancer.

## Data and Code Availability

All data used in this study are available online from public repositories. The Horlings et al. dataset, TCGA-BRCA, METABRIC, and SCAN-B (GSE96058) can each be downloaded directly from their respective sources^31,33,57,59,64,65^. We provide scripts to retrieve and preprocess these open-access datasets, enabling full replication of all analyses presented.

All code is available at https://github.com/Arsuaga-Vazquez-Lab/TAaCGH-Extension and is archived on Dryad^83^.

## Acknowledgments

We want to thank J. Perez-Losada and J.L. Martinez for helpful comments on the manuscript.

## Author contributions statement

R.S. and J.A. conceived the study. R.S., and J.A. designed the methodology. S.J.O.Z., M.P., M.K., G.G.-I., R.S., and J.A. conducted the computational analyses. S.J.O.Z., M.P., M.K., G.G.-I., R.S., and J.A. developed software and analysed the results. G.G.-I. curated the data. R.S. and J.A. supervised the project and acquired funding. S.J.O.Z., R.S., and J.A. wrote the manuscript. All authors reviewed and approved the manuscript.

## Additional information

### Accession codes

Publicly available SCAN-B RNA-seq data used in this study are available through the Gene Expression Omnibus under accession code GSE96058.

