## Supplementary for "*PURPL* identifies a poor outcome subgroup within Luminal A breast cancer"

### Supplementary Material

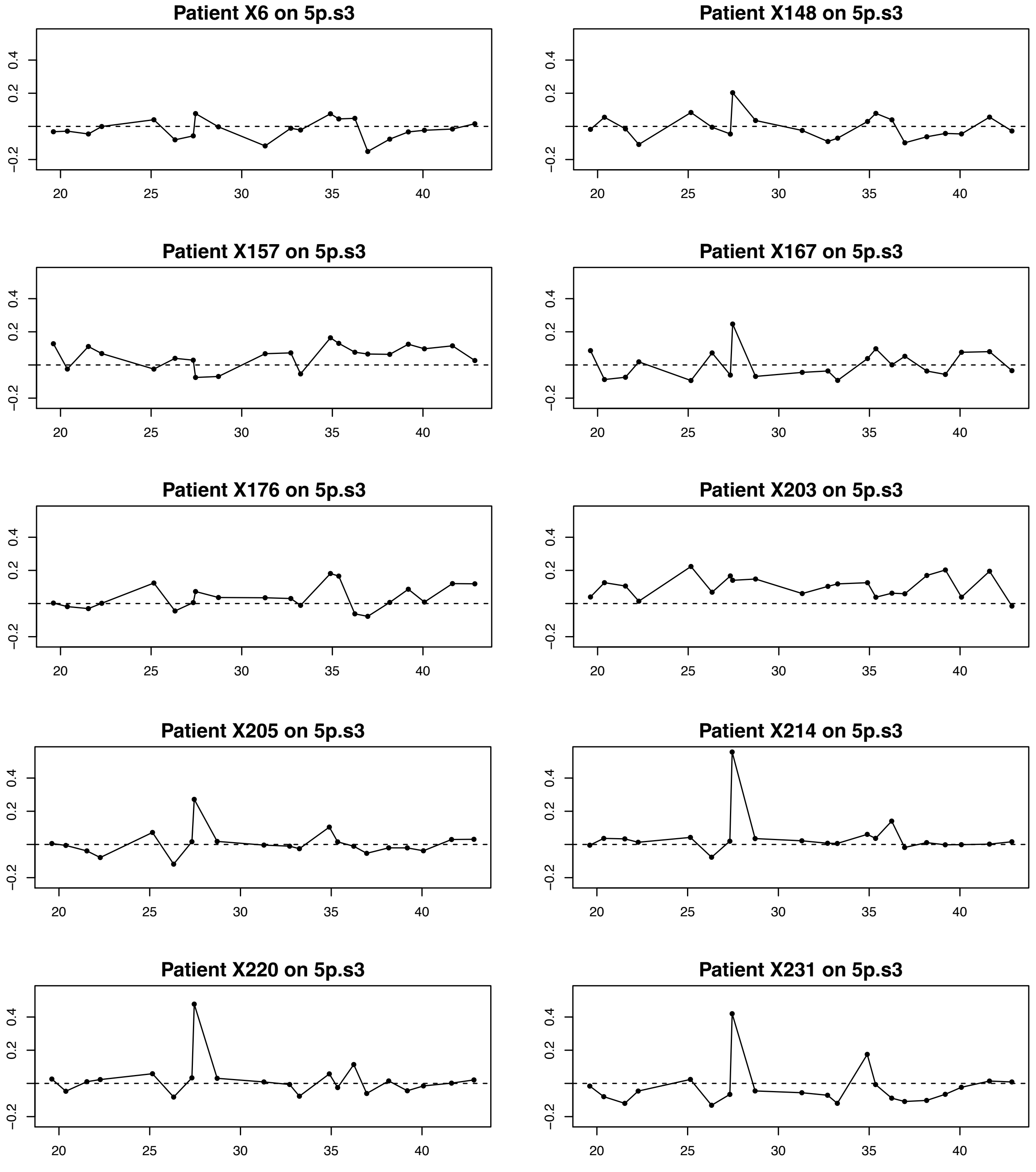


**Figure S1.** Luminal A patient profiles for Horlings dataset on cytobands 5p14.3-p12 in patients X6, X148, X157, X167, X176, X203, X205, X214, X220, and X231. The mean copy number value for Luminal A patients was 0.25 and the mean for the control was 0.11 (with an adjusted p-value=0.015).

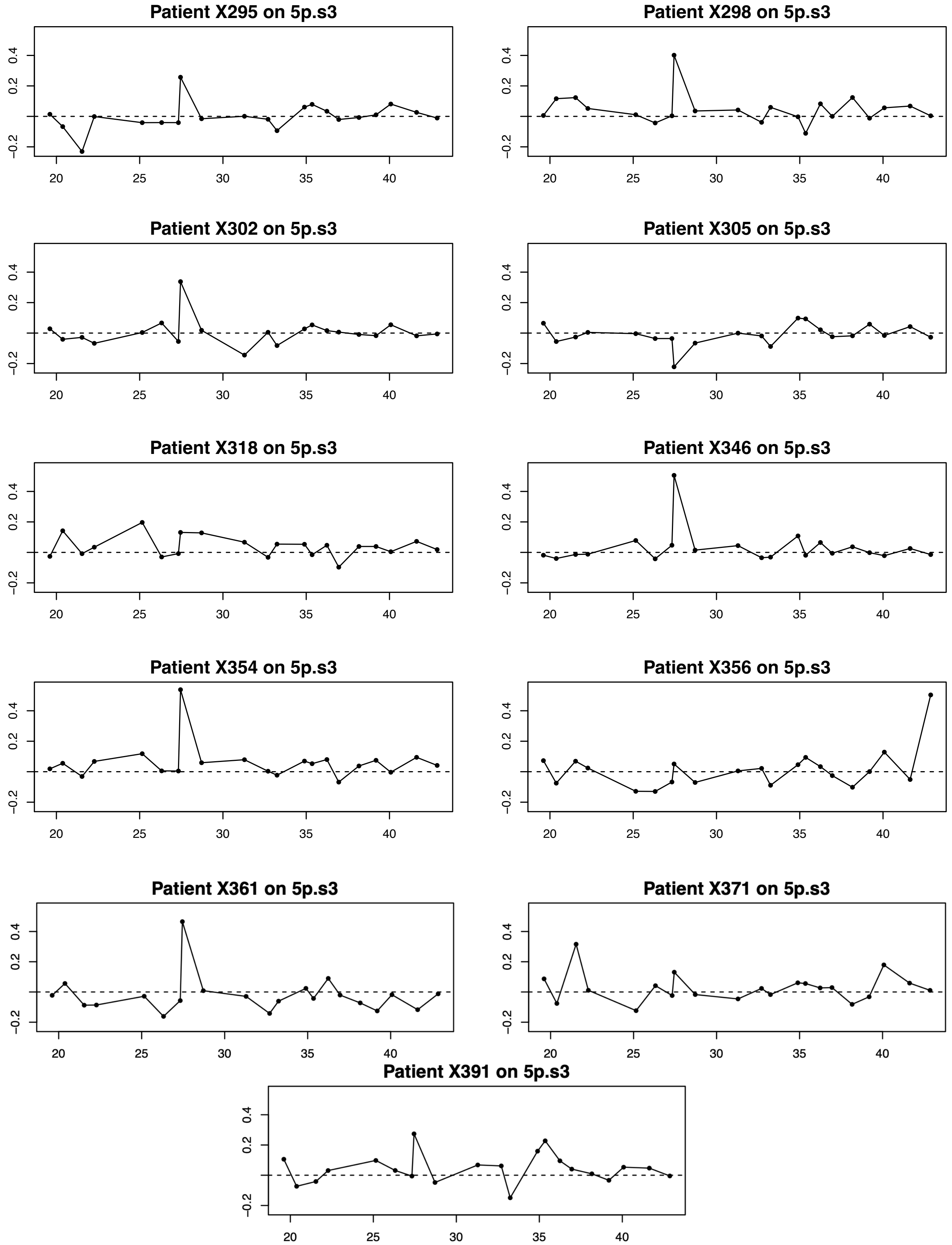


**Figure S2.** (Continued) Luminal A patient profiles for Horlings dataset on cytobands 5p14.3-p12 in patients X295, X298, X302, X305, X318, X346, X354, X356, X361, X371, and X391. The mean copy number value for Luminal A patients was 0.25 and the mean for the control was 0.11 (with an adjusted p-value=0.015)

| ACN | 1 | 2 | 3 | 4 | 5 | 6 | 7 |
|---|---|---|---|---|---|---|---|
| Number of patients | 5 | 331 | 75 | 96 | 17 | 14 | 2 |

**Table S1.** Distribution of Luminal A patients across copy number in the TCGA dataset.

| Gene Status | Patients |
|---|---|
| $TP53_{mut}$ | 49 |
| $TP53_{wt}$ | 484 |
| $MYBBP1A_{mut}$ | 0 |
| $MYBBP1A_{wt}$ | 533 |
| $LYN_{mut}$ | 3 |
| $LYN_{wt}$ | 530 |
| $CCL2_{mut}$ | NA |
| $CCL2_{wt}$ | NA |
| $RBM4_{mut}$ | NA |
| $RBM4_{wt}$ | NA |

**Table S2.** Counts by gene mutation status for TCGA dataset.

| Group | n | Cutoff | M | Lau92 | exactGauss | condMC |
|---|---|---|---|---|---|---|
| $TP53_{wt}$ | 484 | 4.895 | 3.0257 | 0.0540 | 0.0366 | 0.0310 |
| $TP53_{mut}$ | 49 | 4.098 | 1.9448 | 0.5023 | 0.2555 | 0.2508 |
| All Samples | 533 | 4.895 | 2.7602 | 0.1060 | 0.0712 | 0.0693 |

**Table S3.** MaxStat cutpoint summary and associated p-values testing for independence of the Log-Rank statistic $M$ with distinct estimation methods (Lau92, exactGauss, condMC) for TCGA dataset.

| Group | n | Cutoff | M | Lau92 | exactGauss | condMC |
|---|---|---|---|---|---|---|
| $TP53_{wt}$ | 1442 | -2.798 | 2.6043 | 0.0831 | 0.0275 | 0.0332 |
| $TP53_{mut}$ | 167 | -2.478 | 1.3113 | 1.0000 | 0.8284 | 0.8504 |
| All Samples | 1609 | -2.798 | 2.6646 | 0.0906 | 0.0305 | 0.0368 |

**Table S4.** MaxStat cutpoint summary and associated p-values testing for independence of the Log-Rank statistic $M$ with distinct estimation methods (Lau92, exactGauss, condMC) for SCAN-B dataset.

| Group | n | KM corrected (BH) p-value |
|---|---|---|
| $TP53_{wt}$ | 484 | $3.436 \times 10^{-5}$ |
| $TP53_{mut}$ | 49 | 0.0212 |

**Table S5.** Kaplan-Meier corrected (BH) p-values for PURPL gene expression from TCGA dataset.

| Group | n | KM corrected (BH) p-value |
|---|---|---|
| $TP53_{wt}$ | 484 | 0.0002 |
| $TP53_{mut}$ | 49 | 0.0212 |

**Table S6.** Kaplan-Meier corrected (BH) p-values for PURPL expression in TCGA dataset after administrative censoring at 2500 days.

| Group | n | KM corrected (BH) p-value |
|---|---|---|
| $TP53_{wt}$ | 1442 | 0.0182 |
| $TP53_{mut}$ | 167 | 0.1828 |

**Table S7.** Kaplan-Meier corrected (BH) p-values for PURPL gene expression from SCAN-B dataset.

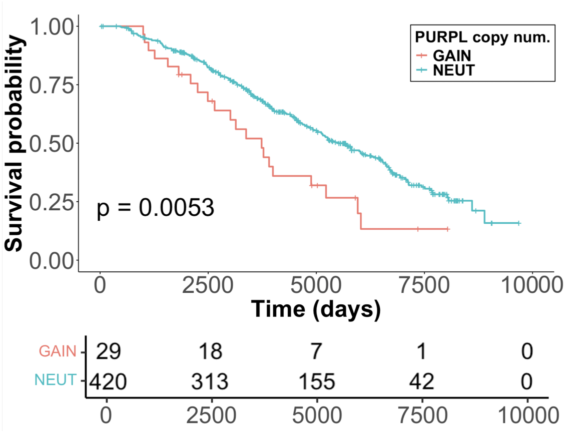


**(a)**

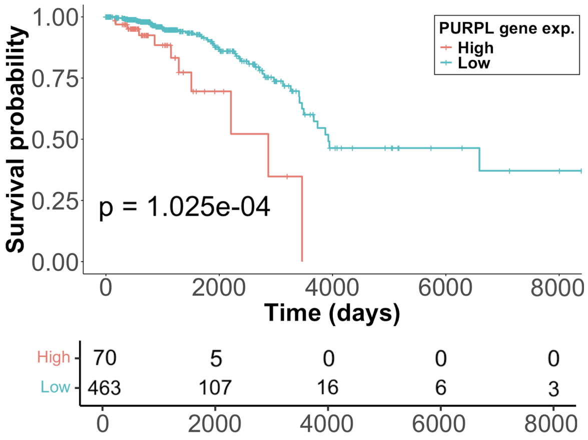


**(b)**

**Figure S3.** *PURPL* copy number and gene expression survival analyses (no administrative censoring): Kaplan-Meier curves stratified by (a) high vs. low *PURPL* ACN in METABRIC and (b) high vs. low *PURPL* expression in TCGA dataset.

| Model | Hazard Ratio | 95% CI | p-value |
|---|---|---|---|
| Standard Cox: *PURPL* only | Not estimable | $[0,\infty)$ | 0.9990 |
| Firth Cox: *PURPL* only | 6.00 | 0.04–112.47 | 0.3545 |
| Firth Cox: *PURPL* + age | 2.71 | 0.02–51.52 | 0.5790 |
| Firth Cox: *PURPL* + node | 5.73 | 0.04–107.63 | 0.3650 |
| Firth Cox: *PURPL* + T stage | 4.34 | 0.03–82.90 | 0.4351 |

**Table S8.** Sensitivity analysis using Firth penalized Cox regression for *PURPL* expression of $TP53_{mut}$ patients in TCGA dataset.

| Model | Posterior HR | 95% CrI | Pr(HR $>$ 1) |
|---|---|---|---|
| Bayesian Cox: *PURPL* only | 0.81 | 0.05–9.84 | 0.4407 |

**Table S9.** Bayesian Cox sensitivity analysis for *PURPL* expression of $TP53_{mut}$ patients in TCGA dataset.
